# Structural Insights into RubisCO Dynamics and Pyrenoid Architecture in *Chlamydomonas reinhardtii*

**DOI:** 10.1101/2025.09.18.677200

**Authors:** Muyuan Chen

## Abstract

RubisCO is the key enzyme that catalyzes the conversion of atmospheric CO2 into organic carbon in photosynthetic organisms. To enhance RubisCO efficiency, eukaryotic algal species pack these enzymes within the pyrenoid, a specialized micro-compartment inside the chloroplast that enriches CO2 for carbon fixation. In the pyrenoid, RubisCO molecules are interconnected by linking peptides and form condensates through liquid-liquid phase separation (LLPS). Using a publicly available CryoET dataset of *Chlamydomonas reinhardtii*, we determine RubisCO structures at 6.5 Å resolution, allowing the trace of the additional densities corresponding to the RubisCO-linking peptides. Through quantitative analysis of RubisCO spatial distribution, we revealed predominant arrangement patterns within the pyrenoid. Additionally, we characterized the structure of pyrenoid minitubules and examined how RubisCOs interact with the membrane network inside the pyrenoid, which facilitates the delivery of substrates for carbon fixation. This work provides valuable structural insights into the LLPS process and enhances our quantitative understanding of carbon fixation within the pyrenoid.

## Introduction

During photosynthesis, energy from sunlight is harvested to convert CO_2_ in the atmosphere into organic carbon. As the key enzyme of carbon fixation and the most abundant enzyme on earth^1^, RubisCO has a slow catalytic rate, as well as relatively poor selectivity of CO_2_ over O_2_ ^2^. To overcome the bottleneck of RubisCO efficiency, photosynthetic species evolved various carbon concentrating mechanisms that actively increase CO_2_ concentration at the site of RubisCO activity. In many algal species, RubsCOs are packed into a distinct micro-compartment inside the chloroplast, called the pyrenoid, which enriches CO_2_ for carbon fixation ^3^.

Inside chloroplast, the pyrenoid is surrounded by a starch sheath, which stores the product of carbon fixation and also separates the RubisCOs from the rest of the chloroplast that surrounds it. Within the starch sheath, hundreds of thousands of RubisCO molecules form a densely packed condensate through liquid-liquid phase separation (LLPS) ^4^. The assembly of RuBisCO condensate inside the pyrenoid is driven by linking peptides, which consists of multiple repeats of RuBisCO-binding motifs connected by intrinsically disordered linker domains ^5^. Those binding motifs form α-helices that are attached to the small subunit of RuBisCO. The linking peptides, such as Essential Pyrenoid Component 1 (EPYC1), are proposed to chain multiple RuBisCO molecules together and facilitate their phase separation ^6^.

To boost RubisCO activity, specialized mechanisms are required to deliver the substrates of carbon fixation into the pyrenoid. In the pyrenoid matrix, thylakoid membranes are remodeled to form an interconnected network of pyrenoid tubules, which penetrate the starch sheath and travel through the RubisCO condensate ^7^. The pyrenoid tubules, which are typically around 100 nm in diameter, are connected to the lumen of thylakoid membranes outside the pyrenoid, and are proposed to deliver CO_2_ in the form of HCO_3_^-^ into the pyrenoid. Inside the pyrenoid tubules, there are even thinner membrane tubules, termed pyrenoid minitubules. Those minitubules connect the pyrenoid matrix to the stroma of the chloroplast ^8^, and are hypothesized to facilitate the delivery of RuBP and 3PGA into the pyrenoid ^9^.

The study of pyrenoids started more than a century ago^3^. The structure of pyrenoid, including the starch layer and pyrenoid tubules, has been visualized by conventional negative stained TEM imaging ^10^, before genetic and immunolabeling studies revealed RubisCO as the major component of the pyrenoid ^11^. Later physiological analyses linked the pyrenoid with carbon concentration mechanisms, showing that the existence of the organelle increases the carbon fixation efficiency in algae. As the research of LLPS advanced, pyrenoid has been identified as a phase separated organelle rather than crystalline or amorphous ^4^. Recent development of cryogenic electron microscopy/tomography (CryoEM/CryoET) makes it possible to directly visualize the pyrenoid at the native condition, and detailed pyrenoid architecture, such as the pyrenoid minitubules, were revealed by CryoET ^8,12^. More recently, EPYC1 was established as the main driving force of the LLPS in pyrenoid, as single particle CryoEM structures suggested that EPYC1 is instrumental in chaining multiple RubisCOs together by binding to RubisCO small subunits ^6,13^.

Recent *in situ* CryoET studies have captured RubisCO structures at subnanometer resolution within the pyrenoid^14,15^. Although multi-model refinement revealed multiple subclasses and potential additional densities associated with RubisCO, the sub-classes do not have sufficient resolution to provide structural information about the extra densities. Furthermore, the high-resolution structures obtained are the result of extensive classification and the exclusion of the majority of particles, meaning that the findings only represent a small subset of RubisCOs in the pyrenoid, making it difficult to provide a complete view of the RubisCO arrangement in the LLPS.

In this study, we reprocessed a publicly available CryoET dataset of *Chlamydomonas reinhardtii* ^14^, focusing on the protein structures and spatial organization within the pyrenoid. Using subtomogram averaging and classification, we determined RubisCO structures with clear densities of the linking peptide. Through per-subunit classification and spatial distribution analysis, we identified the occupancy of linking peptides on RubisCO particles and mapped the predominant arrangement patterns of RubisCOs within the pyrenoid. Finally, we also look into the pyrenoid tubules, examining the minitubules structure inside and exploring how the pyrenoid tubules interact with the RubisCOs in the condensate.

### Structures of RubisCO inside pyrenoid

From the public CryoET dataset of *Chlamydomonas* ^14^, we manually selected 27 tomograms containing the pyrenoid, and automatically identified 157k RubisCO particles using a deep neural network-based particle picker^16^. Through iterative subtomogram refinement with D4 symmetry, we achieved an averaged structure of RubisCO at 6.5 Å resolution using all the particles. The CryoEM map matches well with existing RubisCO structures ^6^, and secondary structure elements are clearly distinguishable (Figure 1, S1).

**Figure 1.**
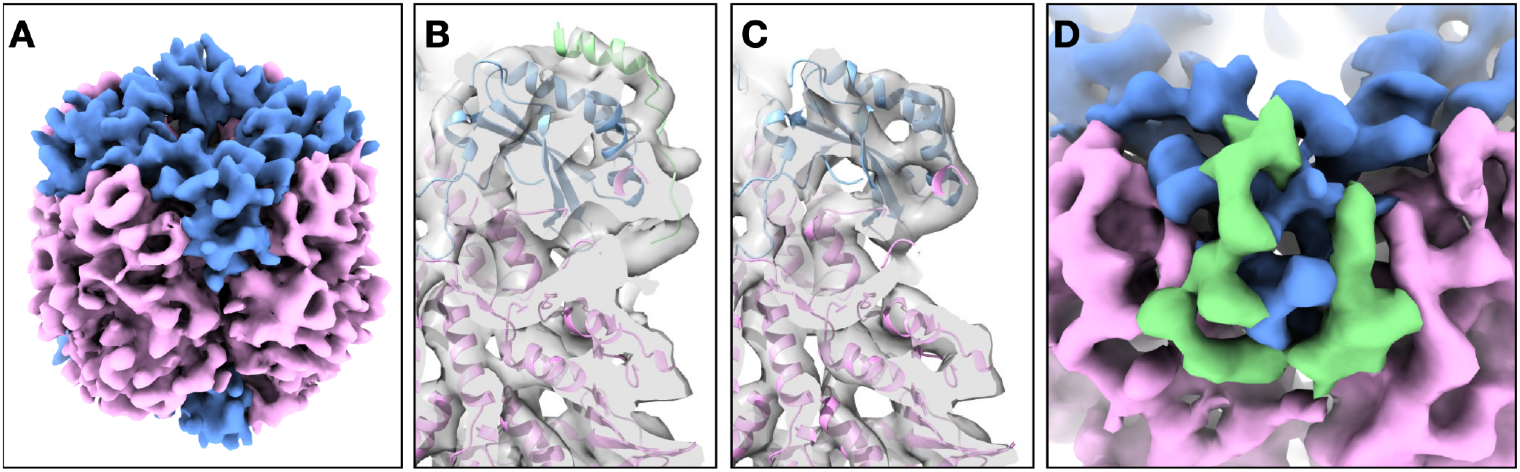
Subtomogram average and classification of pyrenoid RubisCO. (A) Averaged structure of RubisCOs, colored by chains. Pink - large subunit; blue - small subunit. (B-C) Structures from per-subunit classification, with and without the additional linking-peptide density. (D) Zoomed in view of (B) from top, with the additional density colored in green.

We employed multi-model refinement to explore the structure heterogeneity of RubisCO particles^17^. Given that each RubisCO particle comprises eight asymmetrical units, we duplicated each particle eight times based on the D4 symmetry, and performed iterative classification focusing on the one unit. The two classes obtained from this focused refinement displayed distinctly recognizable differences in protein features around the RubisCO small subunit. One class represented the canonical structure of RubisCO, while the other class exhibited pronounced additional densities on the surface of the RubisCO small subunit (Figure 1B-C).

The additional density identified in the second class resembles an alpha-helix-like rod, positioned in the same location as the synthetic fragment of the EPYC1 from the previously reported *in vitro* RubisCO structure ^6^. Given this structural similarity and the fact that EPYC1 is the most abundant RubisCO linking peptide in the pyrenoid ^18^, we hypothesize that the additional density observed in the second class corresponds to EPYC1. It is important to note that, at this resolution, confirming the protein’s identity via sidechain sequence analysis is not feasible ^19^. Nonetheless, for simplicity, we will refer to this additional density as EPYC1 throughout the remainder of the paper.

In comparison to the *in vitro* RubisCO structure that includes only a fragment of EPYC1 ^6^, the structure we obtained from the native pyrenoid reveals several intriguing differences. First, the helical density in the *in situ* structure is tilted slightly towards RubisCO, forming closer interactions with the two helices at the top of the small subunit. Second, the additional density present in the *in situ* RubisCO structure extends further than the helical fragment observed in the *in vitro* structure, traversing the interface between the RubisCO large and small subunits. Compared to the RubisCO-only structure, the additional density in our second class wraps around the small subunit, forming a U-shaped loop. Notably, the U-shaped fold observed in our CryoEM map closely resembles the AlphaFold-predicted structure of EPYC1 ^20^, which fits into the density map with minimal manual adjustments (Figure 1D, S2)

### Arrangement of RubisCO and linking peptides

With the identification of the location and orientation of each RubisCO particle within the tomograms, along with the per-subunit classification of the linking peptide, we can conduct quantitative analyses of RubisCOs arrangement patterns within the pyrenoid.

First, we examined the distribution of linking peptide densities within the same RubisCO particle. Consider two pieces of additional densities on one RubisCO - with the D4 symmetry, there are five possible arrangement patterns (Figure 2A). From our per-subunit classification results, we calculated the conditional probability of each two-EPYC1 arrangement. I.e., the probability of one RubisCO small subunit having the additional density when the other subunit also has the density attached. Notably, the coordinated binding of two adjacent small subunits on the same side of the RubisCO is favored, while all other two-subunit combinations are slightly disfavored. Although the difference in occupancy rates is modest (49% vs. 51%), it is statistically significant (binomial test, p < 0.005 for each class) given the large number of particles analyzed (Figure 2B, S3). The slight preference of adjacent small subunit sites may be attributable to the fold of EPYC1. As the linking peptide loops backward forming the U-shaped structure, the N-terminal end of the loop approaches the binding site of the neighboring RubisCO small subunit, potentially enabling two adjacent RubisCO binding domains of EPYC1 to interact with the two small subunits of the same RubisCO molecule (Figure 2C).

**Figure 2.**
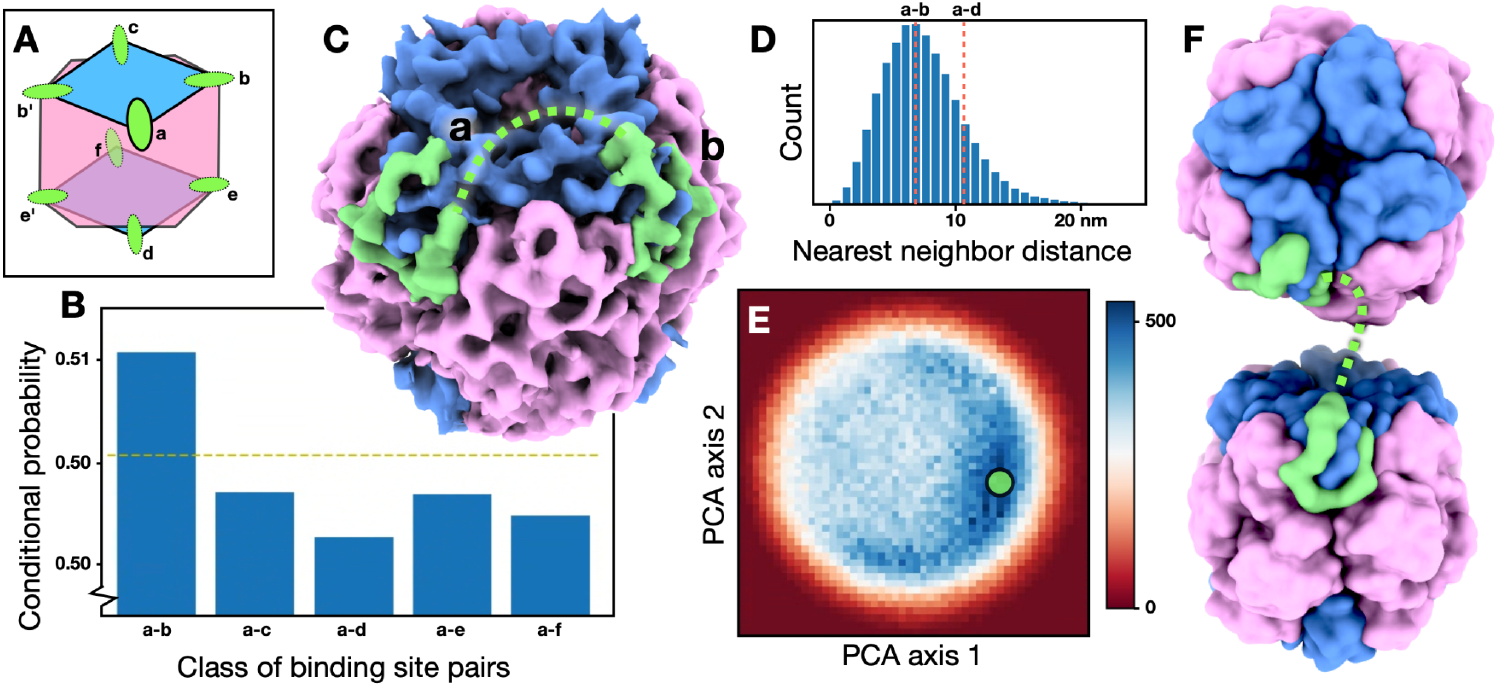
Spatial arrangement of linking peptides on RubisCO. (A) Diagram of RubisCO with possible EPYC1 binding site pairs. (B) Conditional probability of each binding site pair. The yellow dashed line shows the average probability of EPYC1 binding. (C) Subtomogram average of RubisCO class with additional density on two neighboring sites. The green dashed line indicates a possible path of the disordered loop connecting the two sites. (D) Histogram of nearest neighbor distance between adjacent linking peptide densities from different RubisCOs. Red dashed lines show the distance between binding sites from the same RubisCO for comparison. (E) Heatmap of particle pair distribution in the PCA space of relative RubisCO poses. The Green circle indicates the peak location. (F) 3D relative pose of the two RubisCO pairs form the peak location in (E), with dashed green curve showing possible path of connection.

Next, we examined the arrangement of multiple RubisCO molecules in the pyrenoid. In the LLPS system, RubisCO molecules exhibit tight packing, with the center-to-center nearest neighbor distance peaking at 128 Å. Taking the per-subunit classification into consideration, we calculated the nearest neighbor distance between EPYC1 at RubisCO small subunits. When measured from the center of the helical segment, the distance between adjacent EPYC1 densities from two different RubisCO particles peaks at 68 Å (Figure 2D). Notably, this peak is comparable to the distance between two adjacent EPYC1 sites on the same RubisCO particle (69 Å). Furthermore, over 85% of these RubisCO-binding helices have a neighboring binding site within 108 Å, which corresponds to the distance between small subunits of RubisCO on opposite sides. The similarity between the inter- and intra-RubisCO binding patterns suggests EPYC1 does not preferentially bind to the same or different RubisCO molecules when linking the enzymes within the condensate. The binding pattern also allows for simple arrangement patterns where a single EPYC1 alternates between inter- and intra-RubisCO connections, enabling it to chain together three RubisCO molecules using its five helical RubisCO binding domains (Figure S4)^6^.

Finally, we investigated the relative poses between the two nearest RubisCO neighbors. Using principle component analysis (PCA), we mapped the relative poses of neighboring EPYC1-attached RubisCOs into a 2D space, revealing a distinct peak in the heat map of particle distribution. From the center of this peak, we reconstructed particle pairs to obtain an averaged structure comprising two RubisCO molecules. In this structure, the second RubisCO is aligned along the symmetry axis of the first RubisCO, with its own symmetry axis oriented approximately perpendicular to that of the first RubisCO. Notably, in this two-RubisCO arrangement, the N-terminal of one EPYC1 fragment points directly toward the helical region at the C-terminal of the EPYC1 fragment attached to the neighboring RubisCO (Figure 2F, S3).

Although the resolution of the averaged RubisCO-pair structure does not permit tracing of the linking peptide, this geometry suggests a direct connection of EPYC1 between the small subunits of adjacent RubisCO molecules. Additionally, it is important to note that the two-RubisCO geometry is not influenced by the presence of EPYC1 binding. When we conduct the same analysis on particles lacking EPYC1 density, the resulting dominant geometry and its distribution in PCA space remain similar. This observation can be attributed to the dynamic nature of LLPS, in which the linking peptide binds to and unbinds from RubisCOs stochastically ^21^. Thus, even when two adjacent small subunits of RubisCO are not bound by EPYC1 at the moment of freezing, they still maintain a preferred geometry conducive to EPYC1 connection, positioning them for potential future binding events.

### Pyrenoid tubules and minitubules

Pyrenoid tubules are signature structural features of the pyrenoid, traveling through the RubisCO condensate and thought to facilitate the material transfer essential for carbon fixation. Therefore, investigation of the pyrenoid tubules and their interactions with RubisCOs is crucial for understanding the functioning of the pyrenoid ^8,22^. While the overall shape of pyrenoid tubules is highly variable, the minitubules present within them appear surprisingly rigid. Using data from only 773 particles across 11 tomograms, we achieved an averaged structure of pyrenoid minitubule at 28 Å resolution (Figure 3A-B). From the CryoEM map, pyrenoid minitubules take the form of long elliptical tubes measuring approximately 220 x 130 Å in cross-section. Clear separation of lipid membrane leaflets is not detectable on the elliptical tubes, so it is unclear whether the minitubule shell is made of lipid or protein. Inside these tubes, three distinct filamental densities are apparent, each with a diameter of around 30 Å. The number of filaments and their relative positions are consistent across different minitubules, resulting in solid densities in the averaged structure. Along the filaments, a vague repeating pattern of 35 Å is observed. This repeat can be detected through peak searching along the filament in the averaged structure and confirmed by Fourier analysis; however, the resolution of the CryoEM map is insufficient to resolve the individual repeating units of the filament in 3D (Figure S5).

**Figure 3.**
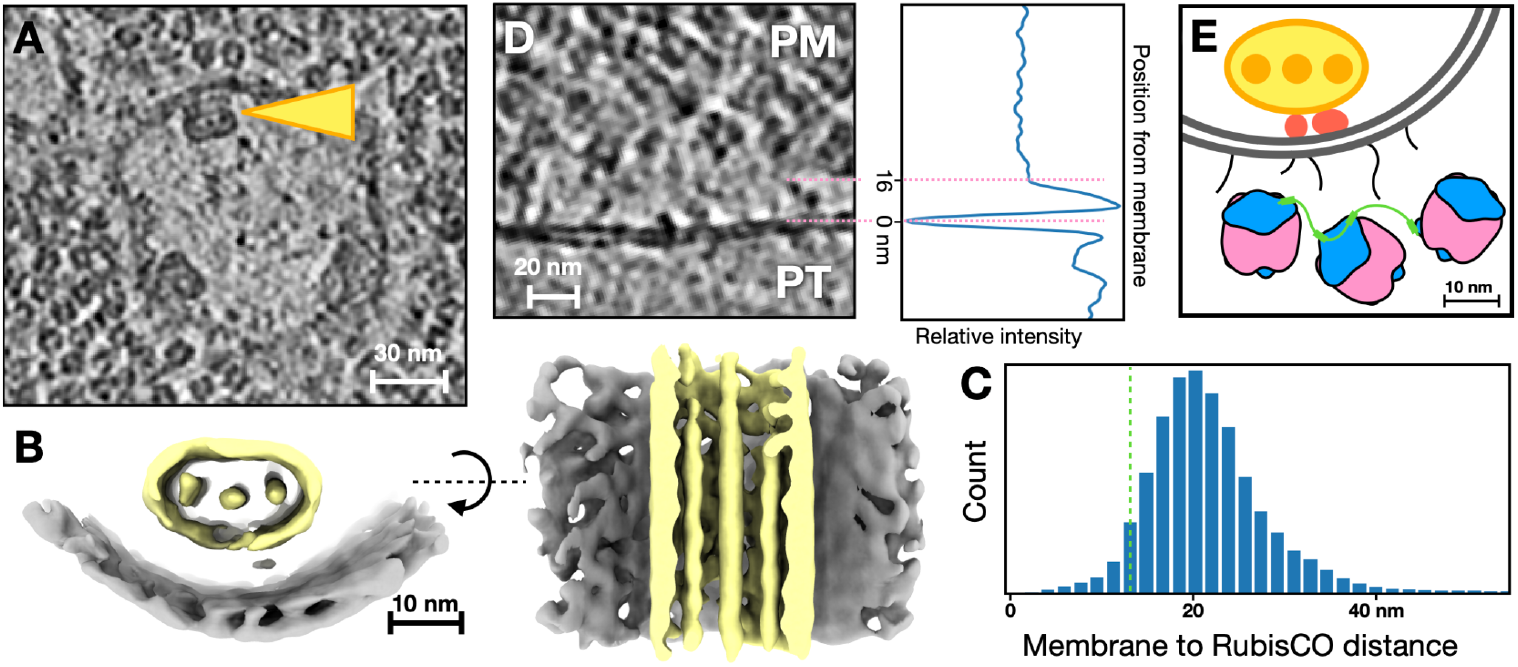
Structure and arrangement of pyrenoid tubules. (A) Slice view of a tomogram, with yellow arrow indicating a pyrenoid minitubule inside the pyrenoid tubule. (B) Subtomogram average of pyrenoid minitubule (yellow) and the adjacent pyrenoid tubule membrane (gray). (C) Histogram showing distance between pyrenoid tubule membrane to the nearest RubisCO. The green dashed line indicates the average distance between two neighboring RubisCOs in the pyrenoid. (D) Polished tomogram slice showing density extending from pyrenoid tubule membranes, repelling RubisCOs. The plot on the right shows the corresponding intensity distribution across the membrane, averaged from multiple subtomograms. (E) Schematic illustration of the spatial relationship between pyrenoid tubules, minitubules and surrounding RubisCOs.

The spatial arrangement of the pyrenoid minitubule in relation to the pyrenoid tubule membranes is also relatively consistent. In the averaged structure, both leaflets of the pyrenoid tubule membrane are distinctly visible. The long axis of the elliptical cross-section of the minitubule is approximately parallel to the nearest pyrenoid tubule membrane, with a distance of 150 Å from the central filament of the minitubule to the nearest pyrenoid tubule membrane.

From the tomograms, connecting densities are occasionally visible between the pyrenoid minitubules and the adjacent pyrenoid tubule membranes. To better visualize their interaction, we conducted classification focused on the region between the membrane of pyrenoid tubules and minitubules. Notably, this classification uncovered a specific class featuring filament-shaped densities that bridge the two membranes. The filament density is tilted at an angle of 30 degrees relative to the direction of the pyrenoid minitubule (Figure S6).

Finally, we examined the interaction between the pyrenoid tubules and the surrounding RubisCOs. The distance from the center of the pyrenoid tubule membranes to the nearest RubisCO center peaks at approximately 200 Å, significantly exceeding the center-to-center distance of 128 Å observed among RubisCOs within the pyrenoid (Figure 3C, T-test p=0.0). This suggests that the pyrenoid tubules may be repelling RubisCO molecules near the membrane. Additionally, aligning and averaging the pyrenoid tubule membranes reveals an exclusion zone of 160 Å around the pyrenoid tubules, which is sufficient to accommodate a single RubisCO molecule (Figure 3D, S7). By refining local tomogram regions using the alignment of surrounding RubisCOs, we achieved a clearer view of the pyrenoid tubule membranes. The polished tomograms show thin, rod-like densities extending from the pyrenoid tubule membrane, which appear to repel nearby RubisCOs. These features suggest the membrane of pyrenoid tubules are covered by membrane proteins for material transport into the pyrenoid. These membrane proteins, and/or the newly arrived material, likely contributes to keeping RubisCO molecules away from the membrane (Figure 3E).

## Discussion

In this study, we determined the structures of RubisCO inside pyrenoid by reprocessing a publicly available CryoET dataset of *Chlamydomonas reinhardtii* ^*14*^. Through subtomogram classification at the per-subunit level, we obtained structures of RubisCO small subunits, both with and without the additional density of linking peptides, hypothesized to be EPYC1. The averaged structure exhibiting the additional density revealed a longer loop compared to the *in vitro* structures of RubisCO-EPYC1, which wraps around the RubisCO small subunit, suggesting a potential interface between RubisCO and the binding protein. Furthermore, our quantitative analysis of the subtomogram refinement results demonstrates the patterns of EPYC1 binding, both within the same RubisCO particle and across multiple RubisCO molecules.

In addition to characterizing RubisCO and its binding peptide, we also examined the structures of pyrenoid minitubules from the same dataset. Contrary to the previous belief that they are merely simple tunnels formed by lipid membranes ^8^, the pyrenoid minitubules exhibit surprisingly rigid structures, comprising three protein filaments encased within a elliptical shell. Given the close interaction between the minitubules and the adjacent pyrenoid tubule membranes, it is plausible that these protein filaments are related to the formation of the pyrenoid tubules and serve as scaffolds that support the membrane network within the pyrenoid.

In comparison to other recent studies of RubisCO structure determination using CryoET, where a majority of selected particles were discarded as “junk” through classification ^14,15^, one advantage of our subtomogram refinement pipeline is that no particle is removed throughout the classification process. Retaining all particles not only resulted in higher-resolution structures, but also preserved the complete dataset for downstream analyses of RubisCO spatial distribution, leading to more accurate assessments of RubisCO arrangement within the pyrenoid. Through per-subunit subtomogram refinement that focuses on protein structure features rather than imaging quality, we generated classes with well-defined structural differences, enabling the modeling of the additional loop into the extra density.

This work also highlights the importance of sharing CryoEM, particularly cellular CryoET data. Unlike studies that focus on single purified proteins, *in situ* CryoET datasets capture the dynamics of thousands of proteins within their cellular context. As a result, reprocessing these datasets can often yield novel insights that go beyond the findings presented in the original publication. We anticipate that, over time, *in situ* structural biology will evolve into a field akin to bioinformatics, where research groups worldwide can make new biological discoveries from publicly available datasets.

Finally, it is important to acknowledge that the structural insights from our work are constrained by the resolution achieved in the averaged structure. Without RubisCO structures with clear sidechain densities, we cannot definitively conclude that the additional density observed on the RubisCO small subunits corresponds to EPYC1. Similarly, without higher-resolution structures of the pyrenoid minitubules, we cannot confidently identify the proteins forming the filaments. These questions will be addressed through future CryoET data collection of the pyrenoid. With a larger dataset at higher magnification, it will become possible to obtain near-atomic resolution structures of RubisCO, revealing a well-defined interface between the enzyme and its binding peptide. Additionally, increased data volume will enable extensive classification, potentially uncovering other RubisCO associated proteins that have lower abundance within the pyrenoid ^5,23,24^.

## Method

### Tomogram reconstruction and particle selection

All tilt series in the public dataset (EMPIAR-11830) were aligned and reconstructed in EMAN2^17^, and the ice thickness of each tomogram is estimated based on contrast distribution along the z axis. The tomograms were sorted by the ice thickness, and 27 thinnest tomograms (<150 nm) with visually identifiable pyrenoid were selected. From the tomograms, 157,929 RubisCO particles were selected using the deep learning based particle picker in EMAN2.

### Subtomogram refinement of RubisCO

Iterative subtomogram and subtilt refinement of RubisCO particles were conducted using EMAN2. An previously published structure of apo state RubisCO, EMD-22401 ^6^, which was lowpass filtered and phase randomized to 20 Å, was used as the initial reference for the refinement. In sum, 7 iterations of refinement for bin2 particles, and 3 iterations for unbinned particles were performed. D4 symmetry was applied throughout the refinement. The Gaussian mixture model based refinement pipeline was used for the alignment of unbinned particles ^25^. The final averaged map has a resolution of 6.5 Å, measured by the “gold-standard” FSC using a soft mask covering a single RubisCO molecule.

For the classification of RubisCO particles, iterative multi-model refinement was first performed for the bin2 particles with D4 symmetry applied. The classification targeted two classes, which showed subtle differences at the tip of RubisCO small subunits. The two classes were then used as initial references for the iterative per-subunit focused classification of unbinned RubisCO particles. In the focused classification, each particle was duplicated eight times according to the D4 symmetry, and a soft spherical mask that covers one RubisCO small subunit and its adjacent large subunits was used to focus the classification. The population ratio between the two classes is roughly 1:1.

To generate subtomogram average of RubisCO with two EPYC1 attachment, we searched through the per-subunit classification results for RubisCO particles with asymmetrical units at the corresponding sites classified as having the additional densities. Notably, we only ensured the target sites to have the EPYC1 density, but not the absence of EPYC1 density on the rest of the sites. That is, for the a-b class in Figure 2C, while all particles in the reconstruction have EPYC1 density at site a and b, the other sites have partial occupancy, so weak EPYC1 density may still be visible. To generate subtomogram averages of two RubisCO pairs (Figure S3B-C), 500 particles closest to the peak in the PCA heatmap were selected, and iterative subtomogram averaging with local realignment was conducted for those particles.

### Modeling of RubisCO with EPYC1

To model the additional densities in the RubisCO subtomogram average, we started from the existing molecular model of purified RubisCO with attached EPYC1 fragments (PDB:7JFO) ^6^, as well as the AlphaFold model of the EPYC1 (AF-Q94ET8) ^20^. The 7JFO model was first fit to the averaged structure in UCSF ChimeraX ^26^, then the corresponding residue (51-72) of the AF-Q94ET8 model was matched to the EPYC1 fragment of 7JFO so it also fits into our *in situ* density map. Afterwards, the AF-Q94ET8 model was combined with the RubisCO structure from 7JFO, and fitted into the subtomogram average using ISOLDE ^27^. Finally, the EPYC1 structure out of the density map was cropped out, leaving the alpha helix and a loop (40-72) surrounding the RubisCO small subunit.

For the modeling of three RubisCOs in Figure S4, RubisCO models (7JFO) were first fitted to the subtomogram averages of 2-RubisCO pairs. Then, helices of EPYC1 RubisCO binding domains from the AlphaFold models were manually placed at their corresponding locations at the RubisCO subunits. Finally, ISOLDE was used to optimize the interface between the full length EPYC1 and RubisCO, as well as the path of disordered loops ^27^.

### Subtomogram refinement of pyrenoid minitubules

773 particles of pyrenoid minitubules were selected from 11 tomograms. Here, we treated the minitubules as filaments and manually traced their curves from the tomograms. The direction of the curves were used as prior for the subtomogram alignment, and the per-tilt translation of nearby RubisCO particles, obtained from the previous subtilt refinement, were used to correct for the local tilt series drifting and improve the averaged minitubule structure. An initial model was directly generated from the particles without a reference, and an averaged structure at 28 Å resolution was obtained through iterative subtomogram refinement. A soft cylindrical mask that covers a 400 Å segment along the tubule was used during the refinement.

To examine the interface between pyrenoid minitubule and pyrenoid tubule membrane, a smaller soft cylindrical mask that covers the region between the pyrenoid tubule and minitubules was used for the focused classification. The iterative focused classification targeted three classes, and resulted in three structures with (A) no density between the tubules, (B) a single filament density between the tubules, and (C) a sheet of multiple filament densities between the tubules (Fig S6).

### Analysis of protein spatial arrangement

Position and orientation of RubisCO particles in tomograms were summarized from the coordinates of particle picking and the results of subtomogram alignment. Distance between RubisCO particles was measured from the center of D4 symmetry, and distance between EPYC1 density was measured from the center of RubisCO binding helix at residue 65 of EPYC1. All spatial and statistical analyses were performed using Scipy ^28^, and the plots were generated using Matplotlib ^29^.

To obtain the dominant poses of two RubisCO molecules, we first posed the central RubisCO particle so it centers at the origin with its D4 symmetry axis aligned with the z axis and the EPYC1 density in the first octant. For each RubisCO subunit with EPYC1, we computed the poses of all of its neighbors with an EPYC1 attachment and converted them into the same coordinate system. So for each EPYC1 pair, there were three 3D coordinates: the center of the second RubisCO, the direction of its symmetry axis, and the location of the RubisCO binding helix of EPYC1. PCA was used to map the 9-value vectors to a 2D space, and a heatmap was computed from the distribution of EPYC1 pairs in the PCA space. Therefore, peaks in the PCA heatmap would represent the preferred relative poses of two RubisCOs in the pyrenoid. In addition to the center of the peak, we also reconstructed particles towards the edge of the peak. All average structures we obtained have two RubisCOs with their symmetry axis pointing perpendicular to each other, and the EPYC1 density of the second RubisCO facing toward the center (Figure S3).

To analyze the spatial arrangement of RubisCO and pyrenoid tubules, we manually annotated the pyrenoid tubule membranes in four tomograms, and calculated the distance from each point along the contour of the pyrenoid tubule membrane to the center of the nearest RubisCO particle. The membrane annotation also provides the normal vector at each point on the membrane, so we extracted 1146 subtomograms at the locations of the membrane, and aligned them by the normal vector (Figure S7). From the averaged image, we computed the intensity distribution across the pyrenoid tubule membrane, and measured the range of the RubisCO exclusion zone. The polished subtomogram in Figure 3D was generated using the subtilt alignment parameters of the RubisCO particles near the target pyrenoid tubules.

---

**Figure S1.**
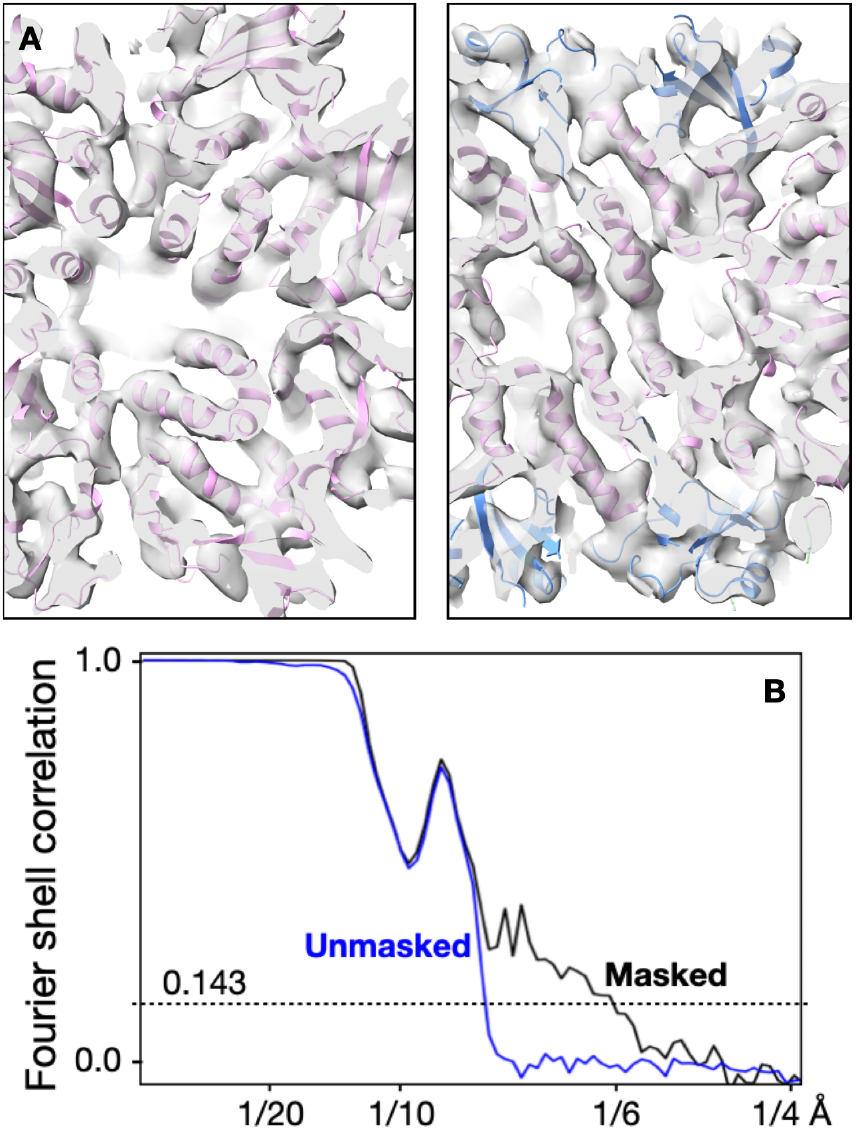
Subtomogram average of RubisCOs. (A) Central slab of the RubisCO structure along the z and y axis, showing densities of secondary structures. (B) “Gold-standard” FSC curve of the RubisCO subtomogram average.

**Figure S2.**
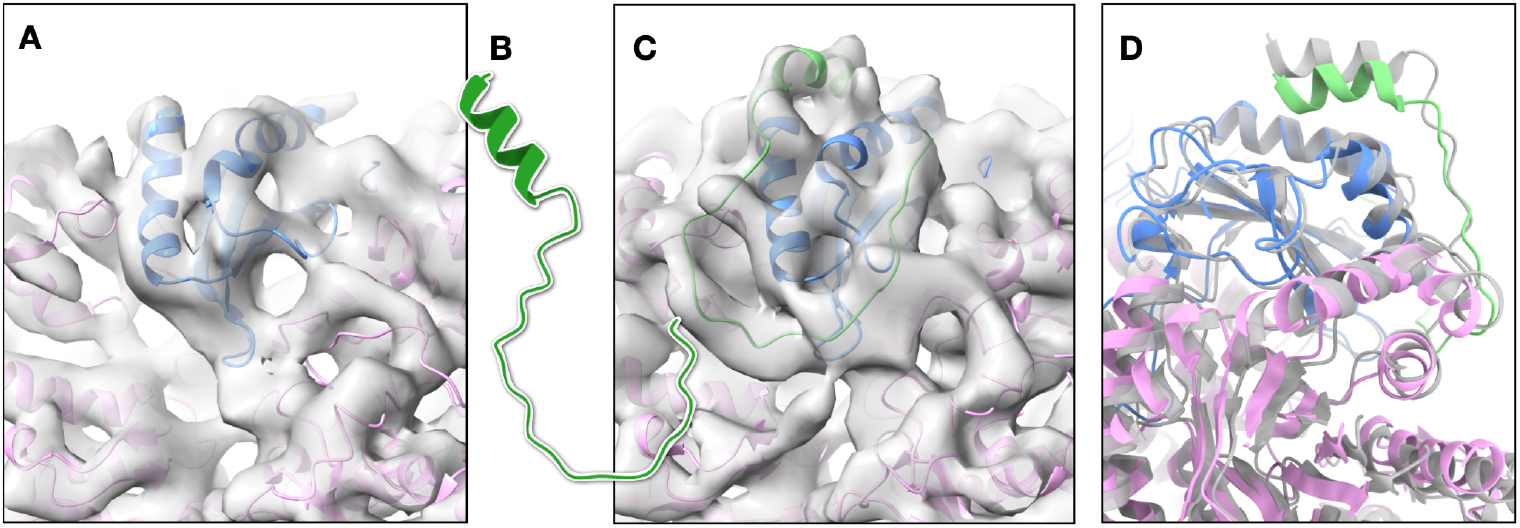
Structure of the linking peptide on RubisCOs. (A) Class of RubisCO structure without the additional density. (B) EPYC1 fragment (residue 40-72) predicted by AlphaFold, showing the U-shaped loop. (C) Class of RubisCO structure with the additional density, and the AlphaFold structure flexibly fitted into the density. (D) Difference between the molecular model fitted to the subtomogram average (pink, blue and green) and the CryoEM structure of purified RubisCO-EPYC1 (PDB:7JFO, gray).

**Figure S3.**
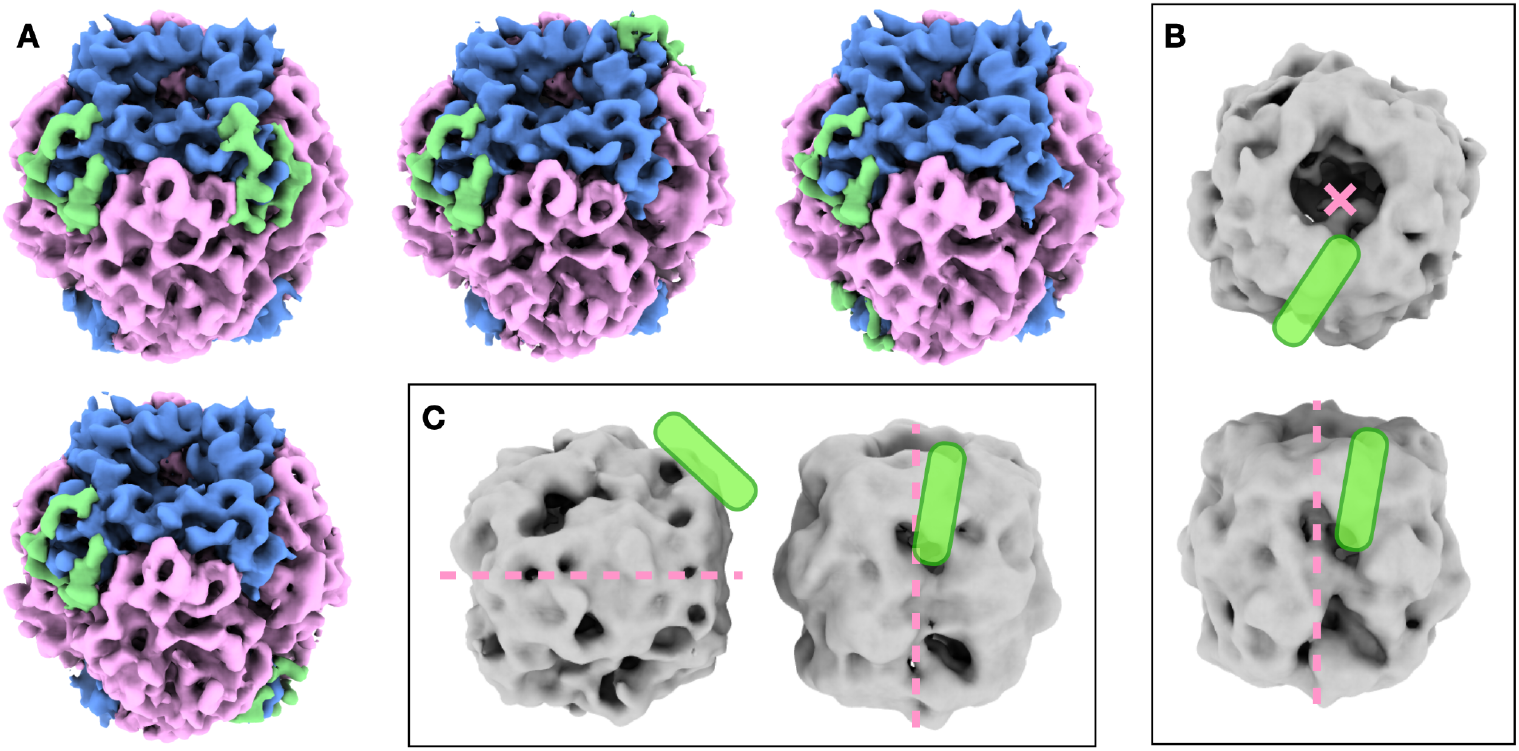
Arrangement of linking peptides on two RubisCOs. (A) Subtomogram averages of RubisCOs with different 2-EPYC1 arrangements. The class a-f is omitted because the f site is blocked by the large subunit. (B) Subtomogram average of two RubisCOs at the peak in the PCA heatmap in Figure 2E. Green rods indicate the positions of the RubisCO binding helices in EPYC1, and the pink dashed line/cross shows the symmetry axis of RubisCO. (C) Subtomogram average of two RubisCOs from particles at the edge of the peak in the PCA heatmap.

**Figure S4.**
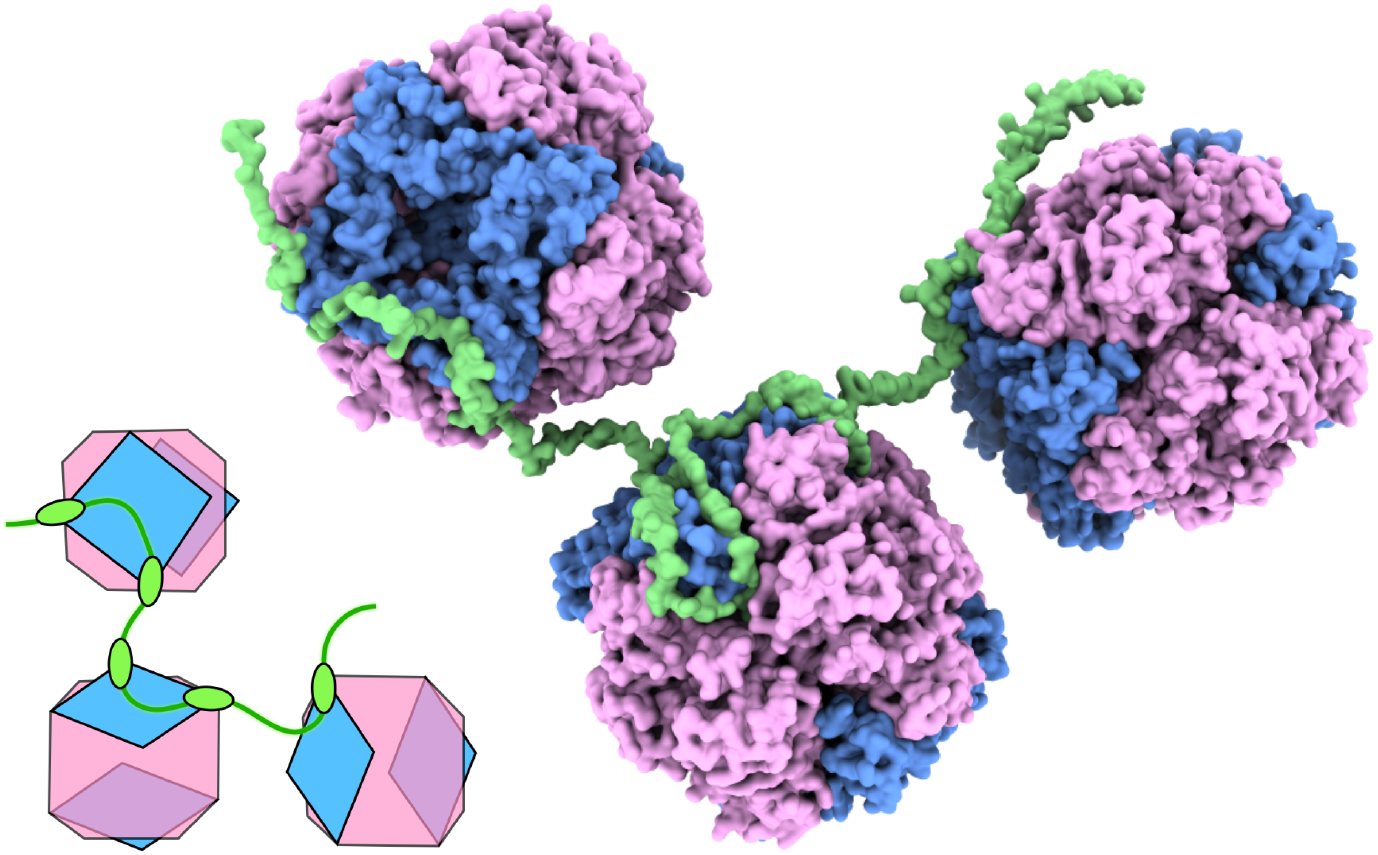
Three-RubisCO arrangement connected by EPYC1. Conceptual diagram and corresponding model of the full length EPYC1 chaining three RubisCOs at their preferred relative poses, by adapting an alternating intra- and inter-RubisCO connection pattern.

**Figure S5.**
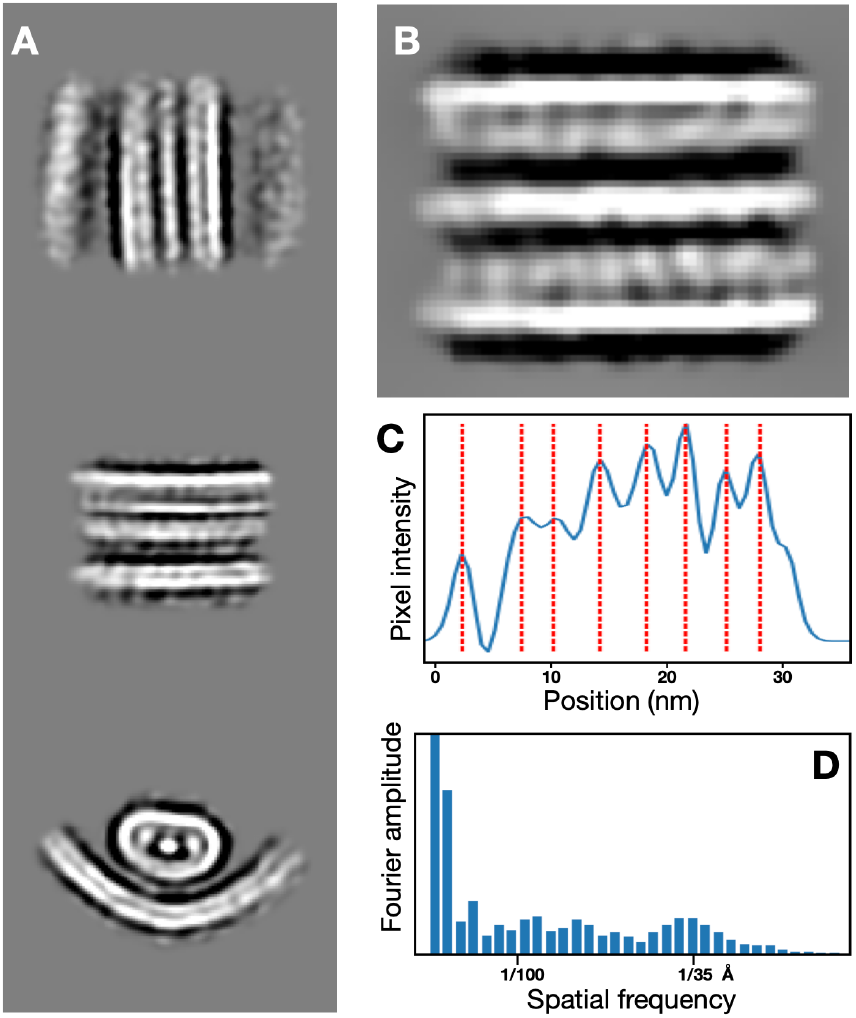
Repeating pattern of pyrenoid minitubules. (A) Orthogonal projections of the pyrenoid minitubule structure. (B) Central slice view of the pyrenoid minitubule. (C) Pixel intensity distribution along the central filament of the minitubule. The average distance between red dashed lines is 36.8 Å. (D) Average of Fourier transform of the three filaments inside pyrenoid minitubule, showing the peak around 35 Å.

**Figure S6.**
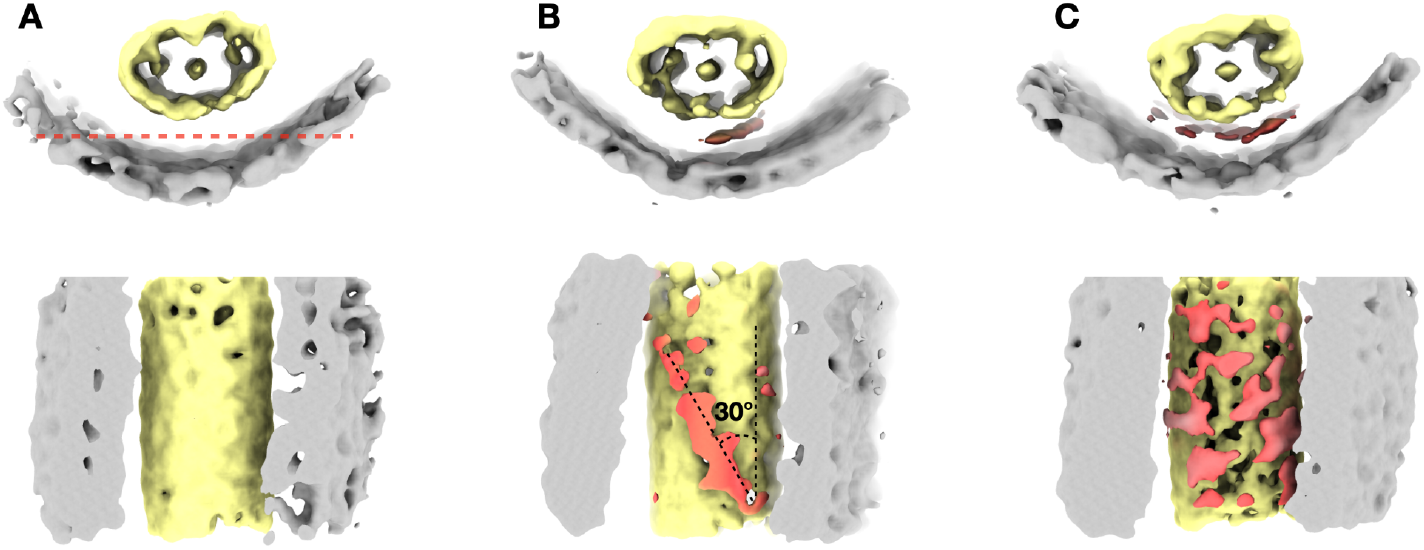
Classification of contacting site between pyrenoid tubule and minitubule. (A-C) Three resulting classes of focused classification. The densities in between the pyrenoid tubule and minitubules are colored in red.

**Figure S7.**
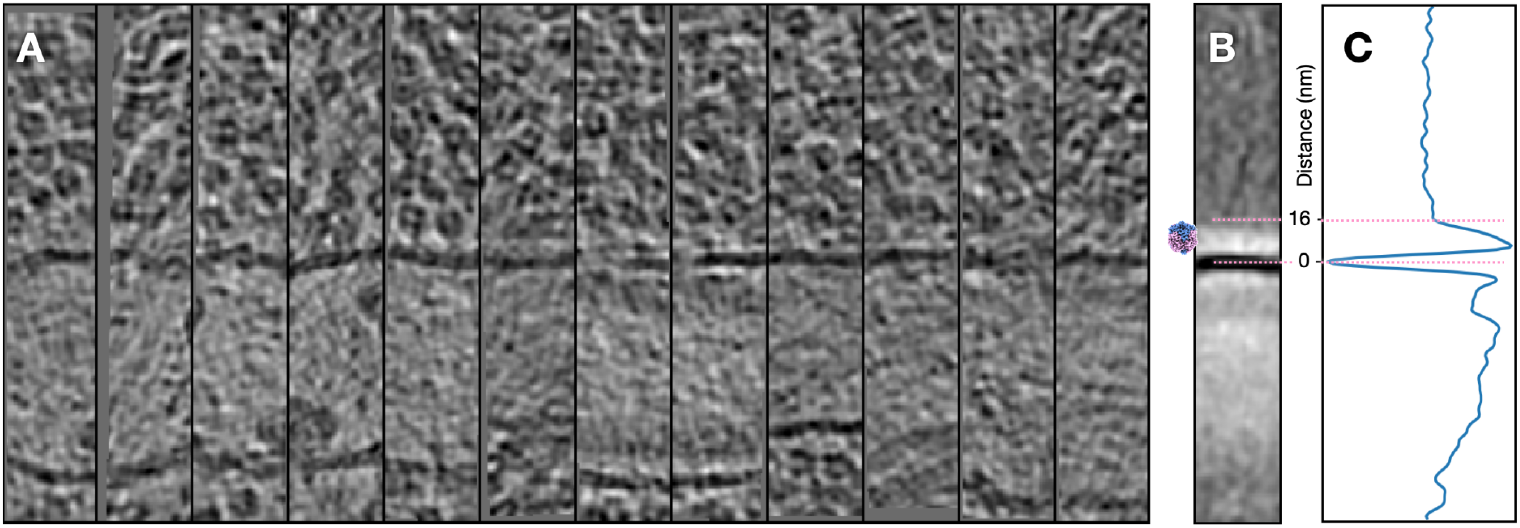
Analysis of RubisCO exclusion zone outside pyrenoid tubule membranes. (A) Examples of aligned pyrenoid tubule membranes. (B) 2D average of pyrenoid tubule membranes. One RubisCO structure was displayed to show the width of the exclusion zone. (C) Averaged intensity of (B), plotted along the y axis.

## Notes

### Competing Interest Statement

The authors have declared no competing interest.

### Funding

- National Institute of General Medical Sciences: R01GM150905

